# Patient Clustering and Classification for Vital Organ Failure Using ICD Code with Graph Attention

**DOI:** 10.1101/2022.11.07.515209

**Authors:** Zhangdaihong Liu, Ying Hu, Gert Mertes, Yang Yang, David A. Clifton

## Abstract

**Objective:** Heart failure, respiratory failure and kidney failure are three severe organ failures (OF) that have high mortalities and are most prevalent in intensive care units. The objective of this work is to offer insights on OF clustering from the aspects of graph neural network and diagnosis history.

**Methods:** This paper proposes a neural network-based pipeline to cluster three types of organ failure patients by incorporating embedding pre-train using an ontology graph of International Classification of Diseases (ICD) codes. We employ an autoencoder-based deep clustering architecture jointly trained with a K-means loss, and a non-linear dimension reduction is performed to obtain patient clusters on the MIMIC-III dataset.

**Results:** The clustering pipeline shows superior performance on a public-domain image dataset. For MIMIC-III, the model gives two distinct clusters that are related to the severity of the diseases. The learnt ICD embeddings present strong power in identifying the OF type in supervised learning.

**Conclusion:** Our proposed pipeline gives stable clusters, however, they do not correspond to the type of OF which indicates these OF share significant hidden characteristics in diagnosis. These clusters can be used to signal possible complications and severity of illness.

**Significance:** We are the first to apply an unsupervised approach to offer insights from a biomedical engineering perspective on these three types of organ failure, and publish the pre-trained embeddings for future transfer learning.

## I. Introduction

ORGAN failure (OF) is the main reason for admitting patients to Intensive Care Units (ICU) and the main cause of death in ICU [1], [32]. The mortality rate remains high for OF patients and is significantly higher for patients with multiple OFs [4]. [3] showed that the most common hospital admission diagnosis of patients that had unplanned transfer to ICU was heart failure (HF) (12%). Moreover, HF is reported to affect over 26 million people globally and has a growing prevalence, especially with an ageing population [30]. The most common diagnosis of unplanned ICU transfers was respiratory failure (RF) (27%). It also has the highest incidence rate in ICU and is associated with high short-term mortality and long ICU stays [28]. A population-based cohort study showed that kidney failure (KF) had the highest one-year mortality rate (18.2%) among all OFs that were investigated [28]. The overall mortality of acute KF is around 20%, rising to over 50% for patients who require dialysis [24].

Patients with these OFs have very poor quality of life, and the cost burden of these OFs results in huge health expenditures for countries [33]. It is crucial for a nation’s health system to better understand the underlying relationships between these OFs so that precautionary measures can be taken and early intervention can be achieved more effectively to improve the mortality and treatment. However, identifying patients with vital organ failure timely and correctly can be challenging due to the broad diagnosis associated with presenting symptoms and variations in patient presentations. Furthermore, the pathological and clinical complexities of those OFs are high.

Electronic health records (EHR) store rich information of patients’ hospital admissions including medical histories, demographics, and symptoms, etc. The International Classification of Diseases (ICD) code is a globally used clinical tool for recording patients’ diagnosis and procedures undergone in hospitals and is widely stored (with little missingness) in most EHR systems. The ninth revision of ICD contains over ten thousand different codes [8]. These diagnostic codes can be collapsed into a smaller number of clinically meaningful concepts to form an ontology. Such structured ontology trees can help us to present more descriptive statistics for easier analysis and interpretation [7]. One popular ontology is created by the Clinical Classifications Software (CCS) [37]. The ICD codes with closer relationships are likely to fall under the same lower-level parent medical concept. In general, and especially for hospital in-patients, they tend to obtain multiple ICD codes during one hospital visit, and an EHR system can track the historical visits of a patient. In [6], they made use of such ICD ontology graph and brought together an attention mechanism and recurrent neural network to predict the onset of disease(s). The ontology structure helped the model to learn the representation of rare conditions and generated predictive patient representations.

There is little research reporting and analysing the clinical characteristics of patients with these OFs such as patient clustering and classification. This paper aims to offer insights into OF patient clustering using diagnosis histories in EHR. We hypothesise that the disease complexities are embedded in the ICD codes assigned to patients during their hospital visits.

In this paper, we adopted the attention mechanism and ICD initialisation approach proposed in [6], in which they showed superiority of applying such mechanism to the ICD ontology in prediction tasks. Additionally, we turned the supervised prediction task into an unsupervised setting for OF patient clustering by employing an auto-encoder (AE) based model architecture. We applied this pipeline to the MIMIC-III dataset on patients with the aforementioned OFs, in order to learn patient groups from the diagnosis histories and having more insights on these OFs from an unsupervised point of view. The experiment pipeline can be divided into three stages: (1) pretrain ICD embeddings from the ontology tree; (2) pre-train an AE embedded with attentions and with layer-wise construction loss only; (3) joint-train the AE with a clustering loss added.

Finally, we modified the above pipeline and carried out a classification task to show that with supervised guidance, the ICD embeddings learnt from ontology can identify different OF patients accurately.

The main contributions of this work include: (1) introducing an OF patient clustering pipeline, where the inputs are ICD embeddings pre-trained with ontology; (2) by adding a nonlinear dimension reduction, this pipeline gives two distinct clusters that are potentially related to disease severity and can be used for complication signalling; (3) we publish the pretrained ICD embeddings^1^ which have strong power in identifying OF types with supervised learning. To our knowledge, we are the first to apply pre-trained ICD embeddings to cluster OF patients. The clustering pipeline composition is novel for this biomedical engineering task. We are also the first to publish the pre-trained ICD embeddings for the convenience of future transfer learning.

## II. Related work

### ICD embedding learning

Learning embeddings for ICD codes, which are typically represented by dense vectors, using machine learning methods has been a popular research topic [15], [23]. The learnt embeddings are often used as features for supervised tasks such as predictions and classifications since they contain rich information for patients’ medical histories. Natural language processing (NLP) techniques are suitable tools to aid the learning because ICD codes are often contained in free text parts of the EHR (e.g. charted clinician notes). [35] used long short-term memory to automatically perform ICD coding given the diagnosis descriptions; [17] combined convolutional neural network and ‘Document to Vector’ to achieve text multi-label classification and automated ICD coding; in recent work [18] used state-of-the-art NLP model BERT to joint learn embeddings for ICD and age to predict diseases.

### ICD ontology and graph neural network

The ICD ontology graph is beneficial for ICD embedding learning. There are over 10,000 codes in the ICD Ninth Revision and significantly more in the later revisions. These codes have clinical hierarchies which can be for better understanding of the relationships between diseases and easier analysis. There are a few widely-accepted ontology schemes such as the one mentioned above (CCS) and SNOMED-CT [25]. Incorporating the ontology tree into ICD embedding learning could enhance the relationships between ICD codes and help to learn embeddings for rare codes. [6] were the first to embed ICD ontology graph into deep neural networks to predict diseases. Later, [34] also employed this ICD ontology graph and updated the attention training to improve the ICD embedding learning. Finally, they used BERT for medication recommendation.

### Deep Clustering

Clustering is an unsupervised method in machine learning, and has been a fundamental tool to learn data structures in an exploratory fashion when no label is given. Since the development of DEC (Deep Embedded Clustering [38]) which combines deep neural network with clustering, deep clustering algorithms have drawn much attention from machine learning researchers. Many deep clustering methods have emerged and they can be categorised into different types based on the loss function. The loss function generally consists of a network loss to learn the latent representations and a clustering loss applied to these representations to achieve the clustering goal [2], [22]. The network loss determines the architecture of the neural network. AE (with reconstruction loss) is the most common architecture for deep clustering models. DEC, DCN (Deep Clustering Network) [39], SR-K-means (Soft Regularized K-means) [10] and K-Autoencoders [45] all adopted this architecture. There are also generative models such as Generative Adversarial Network and Variational AE that are used as the network architecture [11], [46], [47]. There are also methods such as JULE (Joint Unsupervised Learning) [40] and DAC (Deep Adaptive Image Clustering) [5] which are CNN-based and involve only a well-designed clustering loss to extract discriminative features to achieve clustering purpose specifically for images. The clustering can be achieved by using an AE with reconstruction loss only [31]. Those with an additional clustering loss, which is normally added after a pre-train of the network, can simultaneously preserve the structure of the data and achieve clustering. Variations of K-means and KL divergence are most widely applied clustering losses in the discriminative and generative network setting respectively.

Many of the aforementioned methods showed competitive performance compared with supervised tasks on publicdomain datasets such as MNIST [16]. However, we found little work that applies the above techniques to medical applications, and much less to cluster OF patients.

## III. Data

MIMIC-III [12] is a public dataset which contains around 60,000 ICU admissions and over 650,000 diagnoses, recorded using International Classification of Diseases, Ninth Revision (ICD-9). Organ failures are often the main reasons to admit a patient to ICU, which makes MIMIC-III an ideal dataset for our analysis.

To identify the patients with HF, RF and KF, we carefully selected the following ICD codes: all end-level ICD codes under 428 (HF), 518.81 (acute RF), 518.83 (chronic RF), 518.84 (acute and chronic RF), 518.51 (acute RF following trauma and surgery), 518.53 (acute and chronic RF following trauma and surgery), 770.84 (RF of newborn), 584 (acute KF), 669.3 (acute KF following labour and delivery) and 586 (renal failure). In total there are 24 ICD codes. In MIMIC-III, a single patient may have multiple visits (admissions). We included all visits of a patient. Notably, the median and 95th percentile for the number of visits are 2 and 4 respectively.

The data summary is shown in Table I. We selected all patients with one of the three OFs. It is clinically interesting to investigate patients with distinct OFs, and as a consequence of this, the task is simplified. Furthermore, to reduce the randomness/noise in the data, we removed patients with fewer than two hospital visits. This yielded 2216 unique patients in this study, and we named this cohort the *target cohort*.

**TABLE I.**
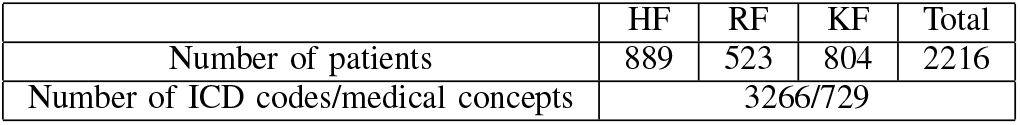
Data summary table. 2216 is the number of patients with only one organ failure and has at least two hospital visits.

|  | HF | RF | KF | Total |
| --- | --- | --- | --- | --- |
| Number of patients | 889 | 523 | 804 | 2216 |
| Number of ICD codes/medical concepts | 3266/729 |  |  |  |

For the ICD ontology, we adopted the scheme created by the Clinical Classifications Software (CCS) [37] which is widely recognised and applied in the literature. There are single-level (ICD codes have only one overall category) and multilevel (ICD codes have hierarchical ancestral categories) CCS categories. We used multi-level CCS to construct the ontology tree. In the ontology tree, the leaf nodes are the billable ICD codes that are actually stored in the EHR system and the upper level ancestors are more general medical concepts. This is illustrated in Fig. 1. For example, ‘Heart Failure’ has three ancestors (apart from the root node which is shared by all ICD codes). Both ‘Heart Failure’ and ‘Atrial Fibrillation’ are end-level ICD codes. They are under a different ‘parent’ node, but belonging to the same ‘grandparent’ node. In this study, there are 3266 unique end-level ICD codes appeared in the diagnoses of the OF patients, and they have 729 ancestor nodes (medical concepts) in the CCS ontology tree.

**Fig. 1.**
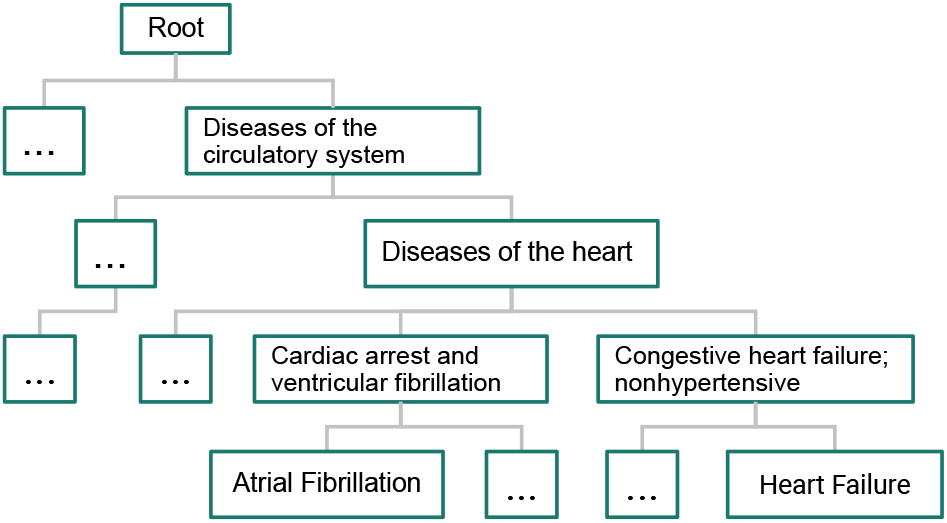
A snapshot of the CCS ontology tree.

## IV. Methods

### A. Analysis pipeline

Fig. 2 A lists the key components from in the clustering pipeline and Fig. 2B shows the details of the end-to-end deep clustering model. The architecture is based around an AE that learns the latent representations of the model input. Clustering is achieved by adding a clustering loss to the bottleneck latents of the AE.

**Fig. 2.**
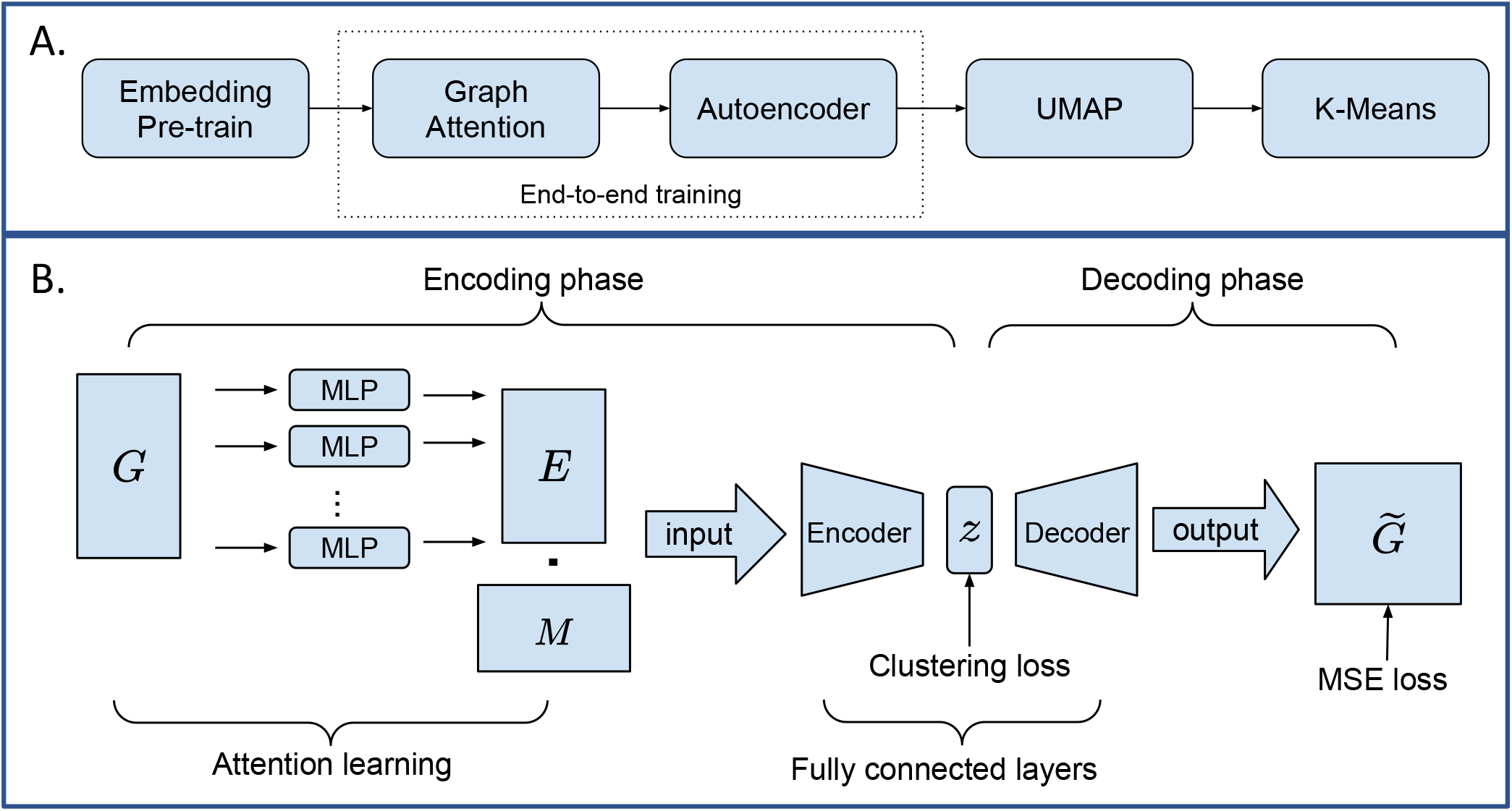
Box A presents the overall clustering pipeline with the key components listed. The dotted box shows the end-to-end training components. Box B shows the details of the end-to-end training. The input of the model is the product of the patients multi-hot ICD encoding matrix *M* and the pre-trained ICD embeddings *G* (trained by GloVe). Each ICD GloVe embedding goes through a multi-layer perceptron (MLP) to learn the attention weights. The concatenation of all MLP outputs is the patient-wise updated ICD embedding matrix *E*. Taking the inner product between *E* and a patient-ICD encoding matrix *M* to serve as the input of a stacked AE. The reconstruction loss is applied between the model input and final reconstruction 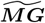 the clustering loss is imposed on the bottleneck latents *z*.

To incorporate ICD ontology into the pipeline, we adopted the approach proposed in GRAM [6]. In brief, [6] imposed an attention mechanism to the ontology tree to establish connections between the leaf ICD nodes and their ancestor medical concepts. The goal is to construct an embedding matrix *E* for the leaf ICD nodes where each embedding in *E* is a weighted linear combination of the ancestor embeddings and the original ICD embedding itself. More specifically, let us assume a randomly initialised embedding matrix *G* containing the initial embedding for all nodes in the ontology tree. The ‘attended’ embedding matrix E contains embeddings for ICD codes only and is constructed by Eqn. 1.

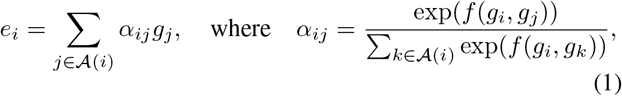

where *e_i_* is the ith column in *E*, corresponding to the *i*th ICD code, *g_i_* the ith column in *G*, representing the ancestors of targeted ICD code, *α_ij_* the attention weights, 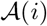 the set for ICD code *i* and all of its ancestors, and *f*(·) in this case represents a two-layer MLP. Taking Fig. 1 as an example again, the ‘attended’ embedding of ‘Heart Failure’ would be a linear combination of the initial embeddings of itself, ‘Congestive hear failure; non-hypertensive’, ‘Diseases of the heart’ and ‘Diseases of the circulatory system’.

The ‘attended’ ICD embedding matrix *E* (concatenation of all MLP outputs in Fig. 2 B) was then further mapped with the patient-diagnosis encoding matrix *M* to serve as the input of a stacked AE.

As shown in Fig. 2 B, the encoding matrix M contains the patients’ diagnosis information. *M* is a *N* × *C* matrix where *N* is the number of patients and *C* is the total number of ICD codes in the dataset. *m_ij_* represents the counts of ICD code *j* from all visits of patient *i*. Fig. 3 illustrates how this map is generated.

**Fig. 3.**
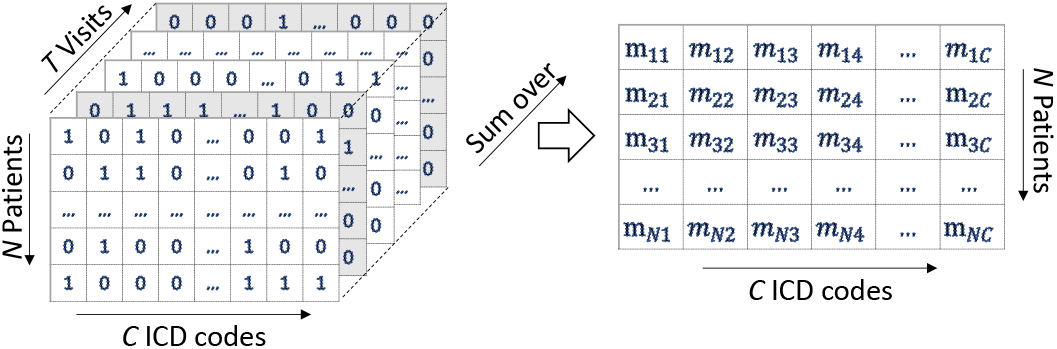
The 3D matrix is a N (number of patients) by C (number of ICD codes) by T (maximum number of visits among all patients) binary matrix indicating whether a patient acquires an icd code in a visit. The visit does not have temporal order, i.e. the *t*th visit for different patients may be at different real-world time. The matrix is padded up to the maximum number of visits using 0 to fill the visits that do not exist for a patient. The multi-hot encoding matrix *M* is the 2D matrix on the right which is summed over the ‘visit’ axis in the 3D matrix.

The initialisation of *G* can be random, however, [6] showed that initialising *G* with GloVe can boost the model performance (details of GloVe initialisation can be found in [29] and [6]). Therefore, we initialised *G* with the same GloVe training, and kept the GloVe embedding dimension as 128. In brief, GloVe is trained on sequences containing all leaf ICD nodes obtained within each visit of a patient and all their corresponding ancestor nodes in the ontology tree. Therefore, GloVe is able to incorporate the hierarchical relationship embedded in the ontology as well as the co-occurrence information between the ICD leaf nodes.

We used layer-wise mean-squared error (MSE) as the reconstruction loss, summing the reconstruction loss between each encoder and decoder layer; K-means loss as the clustering loss and it was applied to the bottleneck latents *z*. The joint loss function is expressed in (2).

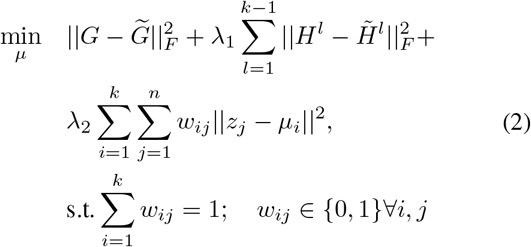

where || · ||_F_ indicates the Frobenius norm, *k* the number of clusters, *H^l^* and 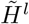 the output of the *l*th encoder and decoder layer respectively, *w_ij_* the binary cluster assignment parameter assigning point *j* to cluster *i*, *z_j_* the *j*th row of the bottleneck latents, and *μ_i_* is the centroid for the ith cluster. The end-to-end training algorithm is described in Appendix Fig. II.3.

We have also considered two other reference pipelines. Details of these pipelines can be found in Appendix I.

### B. Model training and assessment

The training procedure consisted of two pre-trains, a separate ICD embedding pre-train using GloVe (which we name *ICD pre-train)* and an end-to-end *AE pre-train*, and a *joint-train*. Notably, the *AE pre-train* and *joint-trainjoint-train* shared the same end-to-end training pipeline (illustrated in Fig. 2 B) with only loss function different. The *AE pre-train* was trained with MSE as loss only (first two terms in (2)); the *joint-trainjoint-train* has the loss function shown in (2), sum of reconstruction loss and clustering loss. We used Pytorch [27] to train both of the pre-trains and joint-train. The training details can be found in Appendix II.

Regarding the model assessment, we applied the converged model to the dataset, and extracted the bottleneck latents for clustering. Before applying clustering algorithms, we further applied UMAP [21] on the bottleneck latents to reduce the dimensionality to 2. This is to improve the model performance (tested on the public-domain datasets) and convenient visualisation. Then we applied K-means/HDBSCAN [20] to cluster the 2-dimensional UMAP latents. For inference, we investigated the ICD codes as well as their single-level CCS categories belonging to different clusters.

Although we have no ground truth of cluster assignment for patients, it is reasonable to assume the number of clusters is 3 since we are considering three types of organ failure. In this study, we start by setting *k* = 3 for K-means loss and also experimented for *k* equal to 2 and 4. To assess the clustering results, we calculated the Silhouette score [44].

### C. Validation of the deep clustering pipeline on MNIST

To validate the above loss function and training scheme, we tested the deep clustering model (without attention learning) on MNIST [16], an image dataset with 10 classes. The model was trained with the same neural network architecture (AE based), training scheme (a pre-train plus joint-trainjoint-train) and loss function (expressed in (2)). The final clusters are obtained by applying K-means to the 2-dimensional UMAP latents reduced from the bottleneck latents extracted from the converged model. The joint-train algorithm starts from step 11 in Appendix Fig. II.3 in this case.

To assess the clusters, we calculated the standard normalised mutual information (NMI [48]) between cluster outcomes and the true labels and plotted a confusion matrix to visualise the results. Higher NMI indicates better alignment between the clustering outcomes and the true labels.

### D. Classifying OF with ICD embeddings learnt from ontology

To see whether OF patients can be distinguished by their ICD history in a supervised setting, as well as the effect of the attention learning, we carried out a classification task. We set the corresponding organ failure (HF, KF or RF) as the patient label, and turned it into a multi-class classification problem.

We considered three classification pipelines, a main pipeline and two reference pipelines, and they differ only in terms of inputs. The *Main Pipeline* uses the same input as the clustering task, ICD embeddings learnt from the ontology tree (*G* in Fig. 2 B) and patient-ICD encoding matrix (*M* in Fig. 2 B). The first reference pipeline which we call the *No Attention* pipeline uses the product of GloVe embedding *G* and patient-ICD encoding matrix *M* as model input, leaving out the attention training; the *Multi-hot ICD* reference pipeline leaves both the attention training and GloVe ICD pre-train out, using only the multi-hot ICD encoding matrix *M* as input. We show the Main Pipeline in Fig. 4 (two reference pipelines are shown in Appendix Fig. I.2). For the Main Pipeline, the attention learning phase and the initialisation of G are the same as in the clustering pipeline, but the AE architecture was replaced by a MLP with the final layer being three-dimensional and followed by a Softmax activation. During the training, the cross-entropy loss was applied.

**Fig. 4.**
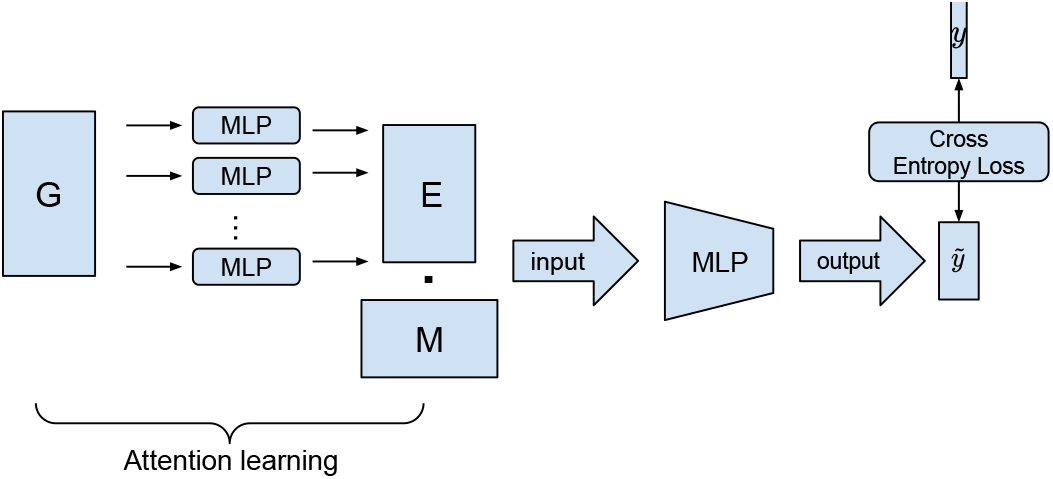
End-to-end OF classification model.

To train the models, we split the data randomly into training, validation and test sets with a ratio of 0.7: 0.15: 0.15. The test set was excluded from the model training and only used for reporting results. We employed a three-layer MLP with layer widths [256, 128, 3], and the training batch size was 200. We calculated the accuracy for all three models to compare their performances.

We also compared these pipelines with three benchmark models: support vector classifier (SVC) [41], random forest classifier (RFC) [42] and XGBoost (XGB) [43]. Since the reference models cannot handle attention nor input samples with different lengths, we averaged the GloVe embedding of all ICD codes for a given patient as input. The data were also split into training and test sets with a ratio of 0.8: 0.2 and with label stratification. We optimised the models with class weights and ran for 10 repetitions.

## V. Results

In this section, we present first the results of testing the clustering model on MNIST, then we move to the main results obtained from MIMIC-III. This part of the results is presented according to the pipeline training order: GloVe ICD embedding training results are shown first; AE pre-train results come second; finally, there are the clustering results and inference obtained from the joint-training. The classification results are presented at the end.

### A. Testing deep clustering model on MNIST

Fig. 5a shows the confusion matrix obtained by applying the pipeline introduced in Section IV-C to MNIST. This result, NMI= 0.929, is at a competitive level with the state-of-the-art clustering models ([9], [10]). Notably, confusion matrices in Fig. 5 show how unsupervised cluster assignments align with the true labels. Since the clusters do not match any specific classes, the blocks are randomly distributed in the matrix. However, if the clusters align well with the classes, one should expect *k* (*k* equals to the number of true classes) distinguished blocks, and distributed in *k* different rows and columns (like shown in Fig. 5a). We also present the result without applying UMAP before cluster acquirement, and found much worse performance (Fig. 5b). This may be explained by the suffering of ‘curse of dimensionality’ of K-means. This is, therefore, the reason why we added UMAP to the analysis of MIMIC-III data.

**Fig. 5.**
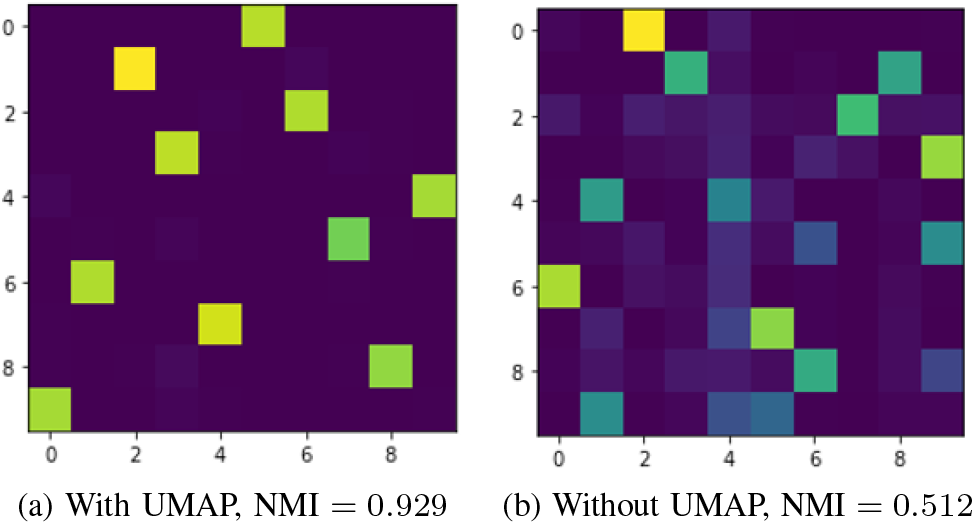
Confusion matrices for clustering results on MNIST with (a) and without (b) UMAP applied. More yellow blocks (relative to green) indicates more instances that fall under the corresponding cluster and class. Note that the numbers on the axis do not indicate the 10-digit classes since this is an unsupervised setting.

### B. OF patients deep clustering on MIMIC-III

#### 1) Interpretation of ICD pre-train

As introduced in Section III, the OF patient cohort comprises over 3266 end-level ICD codes and 729 ancestral medical concepts. GloVe initialisation training gives a dataset-specific embedding matrix for all of the 3995 (3266 + 729) nodes in the ontology tree. It supposes to reveal the co-occurrence information in the OF dataset between the different ICD codes as well as between ICD codes and their ancestors. To be able to interpret the embeddings, we visualised them by applying UMAP to the 128-dimensional GloVe embeddings and reduced the dimensionality to 2. In Fig. 6, we observed clusters where all similar medical concepts gather together such as gynaecological diagnoses (Box B), and diabetes-related diagnoses (Box A). There are also clusters on related diseases such as lung cancer, respiratory diseases and pleurisy in Box D of Fig. 6. Meanwhile, we observed some seemingly unrelated diseases clustered together: skin infection appeared with pulmonary diseases (Box D). The points in Box C are far away from the rest of the points, but they represent a combination of a variety of diseases, e.g., nerve system diseases, nutritional disorders, osteoarthritis and tumour related diseases. The emergence of these clusters might be caused by the co-occurrence of the diseases in this specific dataset or outliers.

**Fig. 6.**
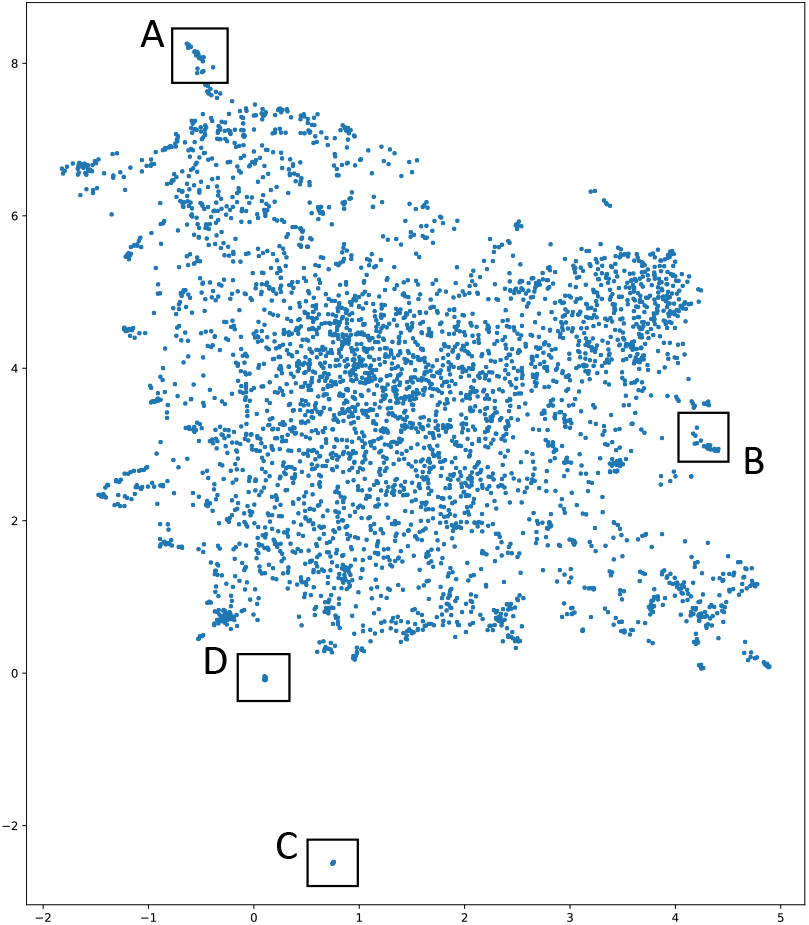
2-dimensional UMAP visualisation of 128-dimensional ICD and medical ancestor embeddings trained by GloVe. Box A contains codes related to medical concepts of diabetes (e.g. Diabetes mellitus with complications) and some other diseases such as ‘Pulmonary heart disease’ and ‘Genitourinary symptoms and ill-defined conditions’. Box B contains all maternal related medical concepts. Box C is a mixture of diseases including ‘Secondary malignancies’, ‘Cancer of bronchus; lung’, ‘Osteoarthritis’, ‘Other hereditary and degenerative nervous system conditions’, ‘Other nutritional; endocrine; and metabolic disorders’. Codes in Box D are ‘Hypertension with complications and secondary hypertension’, ‘Cancer of bronchus; lung’, ‘Skin and subcutaneous tissue infections’, ‘Pleurisy; pneumothorax; pulmonary collapse’, ‘Other lower respiratory disease’ and ‘Inflammatory diseases of female pelvic organs’.

#### 2) AE pre-train

The 128-dimensional GloVe embeddings of all ICD codes and their ancestral concepts were then fed into an AE with reconstruction loss only for pre-train (no clustering loss imposed in Fig. 2 B). Based on the experiments we carried out on MNIST, the visualisation of bottleneck latents after pre-train should be close to the one after joint-train. This statement is also supported by clustering literature [31] where no clustering loss was applied; only AE with reconstruction loss was used for clustering and visualisation. Therefore, visualising the bottleneck latents after pre-train would help us having some ideas on the number of clusters. Since this is a pure unsupervised setting, the selection of the number of clusters should be aligned with the data structure.

As with visualising the GloVe embeddings, we applied UMAP to the bottleneck latents extracted from applying the converged pre-train model to the whole OF cohort. We show this visualisation in Fig. 7a. It looks like the data are moving towards two clusters, and it is possible that the larger cluster may further divide.

**Fig. 7.**
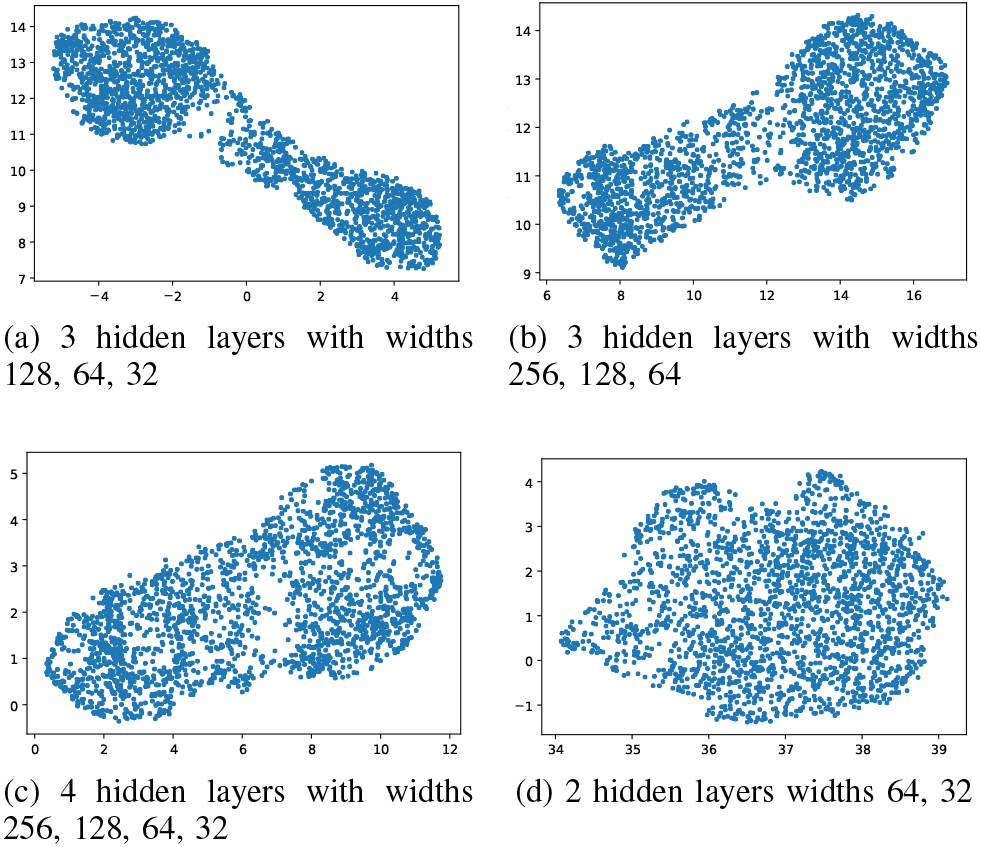
UMAP visualisations for the bottleneck latents after pre-train. The latents are extract from applying the converged model to all of the OF patients. (a), (b) and (c) are trained by the model with 3, 4 and 2 hidden layers receptively, (a) and (b) show a rough representation of two possibly three clusters whereas (c) is hardly showing any structure.

To see the effects of model architecture, we tested with different numbers of hidden layers and layer widths. We observed similar pattern in UMAP visualisation with the same number of hidden layers (with different widths, Figs. 7b) or more layers (Fig. 7c). However, this pattern cannot be learnt with fewer layers (Fig. 7d). This experiment gave us an estimate of the number of clusters to explore during the joint-train. Since Fig. 7a gives the clearest structure and with the simplest model architecture, we used this architecture to carry out the analysis.

#### 3) Clustering results from the join-train stage

We ran the joint-train stage for several repetitions of each *k* ∈ {2, 3, 4}. The clustering visualisations for all *ks* look very stable (Fig. 8). All cases in Fig. 8 display two distinguished clusters. We also observed that as the training proceeds, the longer-shaped cluster tend to be stretched out even further, with a long tail (an example is shown in Appendix III) and no further distinguished division appeared. This kind of visualisation might be a sign of over-training. Therefore, we stopped the training before the cluster became too stretched out. We then moved to interpreting the two clusters presented in Fig. 8.

**Fig. 8.**
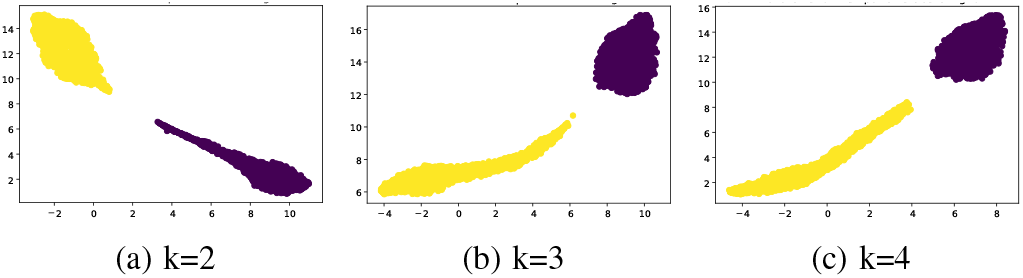
UMAP visualisations for the bottleneck latents after joint-train. The latents are extract from applying the converged model to all of the OF patients. (a), (b) and (c) are trained with 2, 3 and 4 clusters (*k* = 2, 3 and 4) respectively. The two colours are clustering labels assigned by HDBSCAN.

The Silhouette scores for the three sets of clustering results shown in Fig. 8 are 0.800, 0.748 and 0.740 for *k* = 2, *k* = 3 and *k* = 4 respectively. We focused on the *k* = 2 case since it has the highest Silhouette score. Moreover, all *ks* gave very similar results (results for *k* = 3 and 4 can be found in Appendix IV). We first investigated the ICD codes in the two clusters. Due to the large number of ICD codes involved, we applied CCS single-level category to represent the ICD codes to aid interpretation and visualisation. We visualised the CCS single-level category by a range of percentiles based on occurrence frequencies, and focused on interpreting the CCS categories with the top 10-20% occurrence. This is due to the fact that the most occurring ICD codes/CCS categories are generally diseases with high prevalence in the population which is not helpful in distinguishing the clusters. We present the results for the top 10% CCS categories and top 20% most-occurring ICD codes (split into 4 ranges) in Appendix IV.

The top 10-20% CCS categories are shown in Fig. 9 (the same figures for the cases of *k* = 3 and = 4 are shown in Appendix IV). The most commonly occurring CCS category in Fig. 9 is ‘Chronic ulcer of skin’ which can be a complication for all OFs, and it only appears in cluster 1 for this range. We also observed other OF related CCS categories belonging only to cluster 1 in this range such as ‘Heart valve disorders’, ‘Urinary tract infections’, ‘Pneumonia’ and ‘COPD’ which are related to HF, KF and RF respectively. The unique and OF-related categories belonging to cluster 2 in this range include ‘Acute myocardial infarction’, ‘Chronic kidney disease’, ‘Other upper respiratory disease’ which can also be related to HF, KF and RF, respectively. Therefore, the clusters are not grouped by failing organs, but by severity of the diseases in some way: the occurrence frequency for cluster 1 is larger than cluster 2 (normalised by cluster sizes), i.e. the patients in cluster 1 have more diagnoses than those in cluster 2. Moreover, some diagnoses that are unique to cluster 1, such as ‘Chronic ulcer of skin’, ‘Coagulation and hemorrhagic disorders’ and ‘Secondary malignancies’, are signs of more severe deterioration in patients with organ failure [14], [26].

**Fig. 9.**
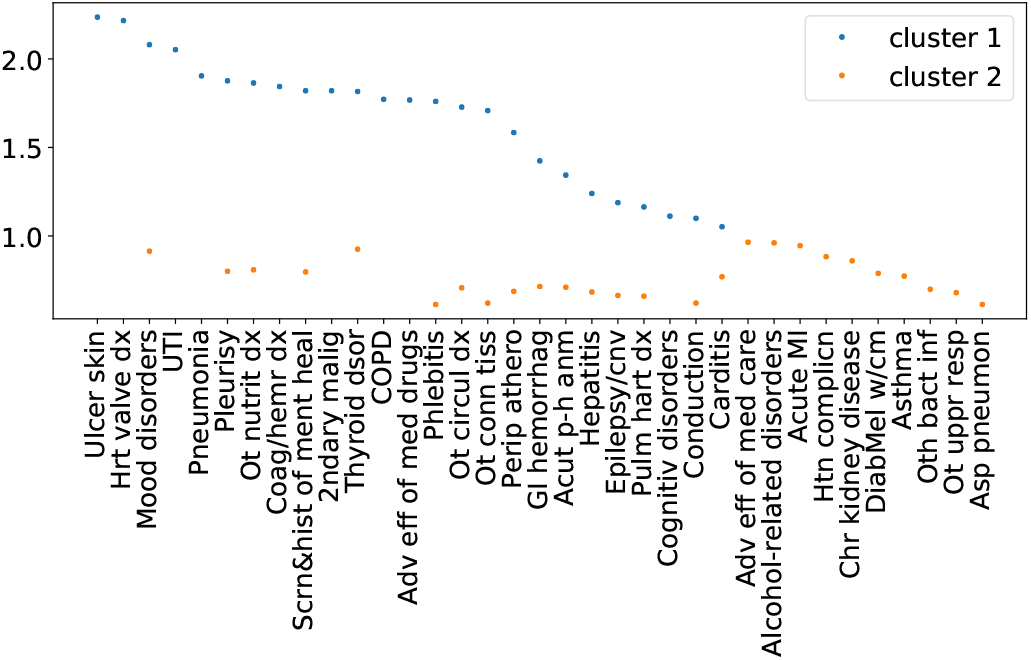
CCS single-level categories with the top 10 to 20 percent occurrence in the two clusters. The occurrence is normalised by cluster size and ordered in decreasing orders. The label ticks are abbreviated category names. The full category names can be found in Appendix V.

### C. Comparison with DCN

We compared our work with the widely applied deep clustering method Deep Clustering Network (DCN) [39] which also has an AE architecture and is extensively used as a benchmark model. DCN also has two loss terms: a reconstruction loss and a K-means loss. We used the same hyper-parameters as in the original where we can. However, DCN does not make use of any attention mechanism. Therefore, we used the average of the code embeddings as input. We ran DCN for *k* = 2 and *k* = 3 (in K-means), and found that although it also displayed a pattern of two clusters as shown in Fig. 10, the stability between different ks and different runs is much worse compared with our method.

**Fig. 10.**
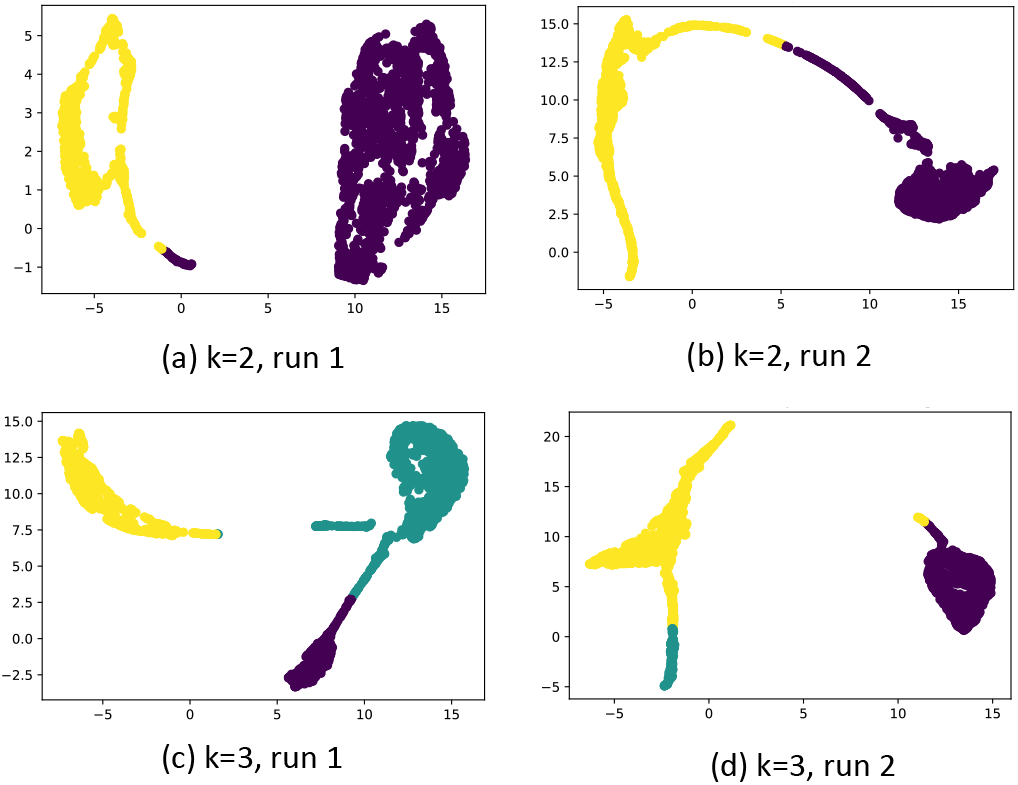
UMAP visualisation for DCN setting *k* = 2 (subplots (a) and (b)) and *k* = 3 (subplots (c) and (d)). We ran each setting two times. The colors are predicted labels given by DCN.

DCN [39] did the same MNIST task as we presented in Section V-A and the NMI reported in [39] (0.81) is significantly lower than using our proposed method (0.93).

### D. Results for classification

For the supervised task, we classified the OF labels using the ‘attended’ ICD embeddings, and compared it with the two reference models, without attention (No *attention)* and using only the patient-ICD encoding matrix *(Multi-hot ICD)* as input features and three benchmark models, SVC, RFC and XGB. As shown in Table II, the main model was able to classify the type of OF significantly better than all of the reference and benchmark models we considered.

**TABLE II.** OF classification results and input dimension for the main model, two reference pipelines, *No attention* and *Multi-hot ICD*, and three benchmark models, SVC, RFC and XGB. Note*: there is no randomness involved in XGB (since we did not use sub-sampling) and SVC.

|  | Main model | No attention | Multi-hot ICD | SVC* | RFC | XGB* |
| --- | --- | --- | --- | --- | --- | --- |
| Input Dimension | 128 | 128 | 3266 | 128 | 128 | 128 |
| Mean accuracy (std) (%) | 99.28 (0.46) | 90.50 (0.62) | 95.81 (0.61) | 90.32 | 78.72 (0.86) | 84.01 |

## VI. Discussions

The identification, through clustering and classification, of heart failure, respiratory failure and kidney failure patients has been a challenge in both machine learning and clinical practice.

We combined the best of several published works such as using layer-wise reconstruction loss, two-stage training and adding graph attention. At the same time, we also tried to make the architecture stay low complexity such as using the discriminative AE plus K-means architecture. Therefore, the model training does not require extensive computation power. Moreover, we explored different clustering loss functions such as adding regularisation terms and replacing K-means loss with fuzzy c-means loss, however, these modifications did not bring extra gain in NMI or Silhouette scores. Therefore, we carried out this clustering pipeline and added the attention mechanism to cluster the OF patients in an end-to-end training fashion.

To investigate the impact of input features on the clusters, we further compared the following scenarios: 1) feeding the product of E and M in Fig. 2 B directly to K-means; 2) applying UMAP to the product of E and M, and then feeding the reduced features to K-means; 3) applying K-means directly to the bottleneck latents without UMAP. We set *k* = 2 for all the above scenarios. The Silhouette scores are 0.5267, 0.5739 and 0.5685, respectively, for the three cases, which are significantly lower than our proposed pipeline (Silhouette score = 0.8005).

Compared with the previous deep clustering works which were mostly proposed in other areas such as computer vision, our proposed pipeline is designated for application in clinical settings - in particular, we extended the encoder in the AE to integrate the diagnosis codes and their ontology using graph attention to learn more stable representations. As a potential consequence, we observed better stability of our pipeline compared with a stereotypical benchmark model, DCN in the clinical task. For the common MNIST task which has true labels and was implemented in almost all deep clustering works. Our pipeline gives higher NMI than other deep clustering methods considered in this work ([10], [38], [45], [46]). We attribute this partially to our application of the non-linear dimension reduction method, UMAP.

Overall, this paper presents a pure exploratory analysis, and the unsupervised setting poses several challenges to the analysis which are common issues of unsupervised learning. First of all, it made the interpretation of the results challenging. Since we had no ground truth, it is difficult to choose the assessment measure and inference approach. Apart from the Silhouette score, we investigated the ICD codes occurring most frequently in different clusters. However, we are aware that there are other ways to interpret the clusters and we may get different information by investigating different measures. Secondly, the clustering task made it hard to assess the model. By embedding the graph attention into the end-to-end clustering pipeline, it makes the comparison with other benchmark methods (such as K-means and HDBSCAN) difficult since this would require the benchmark methods being able to handle graph attention as well. To demonstrate the efficacy of the clustering method, we tested the clustering part of the pipeline (without attention) on MNIST. We acknowledge that this is not a perfect validation, but this is a representative task given the popularity of MINIST in computer vision and signal processing. We further compared a few different sets of input features which were all inferior to our proposed set. We also studied the stability of our current results including exploring different analysis pipelines, AE architectures (number of layers and layer widths) and assigning different *ks* to K-means. Our results showed fair robustness, nonetheless, other hyperparameters can be tuned which may give different results. Moreover, this work only considered patients recorded with only one organ failure. It can be served as a starting point for studies that include patients with multiple OFs in real-world scenarios. One other limitation of this work is that it only uses ICD information; integrating other data modalities such as demographics, procedures and vital signs will be valuable future work.

## VII. Conclusions

This paper proposes an unsupervised learning pipeline for clustering OF patients in MIMIC-III using ICD diagnosis code which has been rarely studied so far, via graph ontology learning and deep neural networks.

We tested this deep clustering model on the public-domain dataset, MNIST, and found that if we add UMAP, a nonlinear dimension reduction method, to the bottleneck latents before clustering, the model performance can be improved significantly. It achieved an NMI of 0.929 – a competitive level of performance with the state-of-the-art clustering algorithms even without the use of convolutional layers.

We discovered two clusters for the OF patients from the model. This discovery is stable to the AE architecture when there are enough hidden layers (3 in this case) to learn such structure, and is robust to the number of clusters to which we assigned K-means during the joint-train learning. The clusters produced by the model did not correspond to the three OFs well. Instead, the two clusters rather related to the severity of the patients, one group having considerably more diagnoses than the other and focusing on different sets of diseases. This outcome may suggest that these three OFs are not separable only based on the ICD information; these groups of patients share similar underlying characteristics or the complexity of the underlying structure is too high to be learnt in this way. However, this model can potentially be used in clinics as a severity identification tool for patients with these OFs and to flag the possible complications that may arise from organ failure.

Furthermore, we carried out the classification analysis and proved that the ‘attended’ ICD embeddings have superior performance in classifying the OF labels which exceeded the two reference and three benchmark models we considered. To our knowledge, we are the first to use GloVe embeddings to perform clustering and classification tasks on heart failure, respiratory failure and kidney failure patients.

## Acknowledgment

Z. Liu and G. Mertes would like to thank the Suzhou Industrial Park for the funding support; Z. Liu was supported by the Jiangsu Provincial Double Innovation Talent Programme; Y. Hu was funded by the Shanghai Municipal Health Commission General Project (202040083); D. A. Clifton and Y. Yang were funded by the EPSRC (EP/N020774/1); this work was supported in part by the National Institute for Health Research (NIHR) Oxford Biomedical Research Centre (BRC), and in part by an InnoHK Project at the Hong Kong Centre for Cerebro-cardiovascular Health Engineering (COCHE). DAC is an Investigator in the Pandemic Sciences Institute, University of Oxford, Oxford, UK. The views expressed are those of the authors and not necessarily those of the NHS, the NIHR, the Department of Health, InnoHK – ITC, or the University of Oxford.

## Appendix I

### Alternative Clustering Pipelines and Reference Classification Pipelines

In addition to the pipeline that was introduced in Section IVA, we also experimented on two other clustering pipelines to compare the findings. Running the pipeline in Fig. I.1a helped us to simplify the experiment and find better activation functions to avoid problems such as vanishing gradients. However, by running this pipeline, we did not obtain distinguishable clusters from the visualisation. The pipeline in Fig. I.1b was our initial main pipeline, however, it gave very similar results to our current main pipeline and ran slower due to a more complicated structure. Therefore, we discontinued this pipeline.

Fig. I.2 illustrates the reference classification pipelines.

**Fig. I.1.**
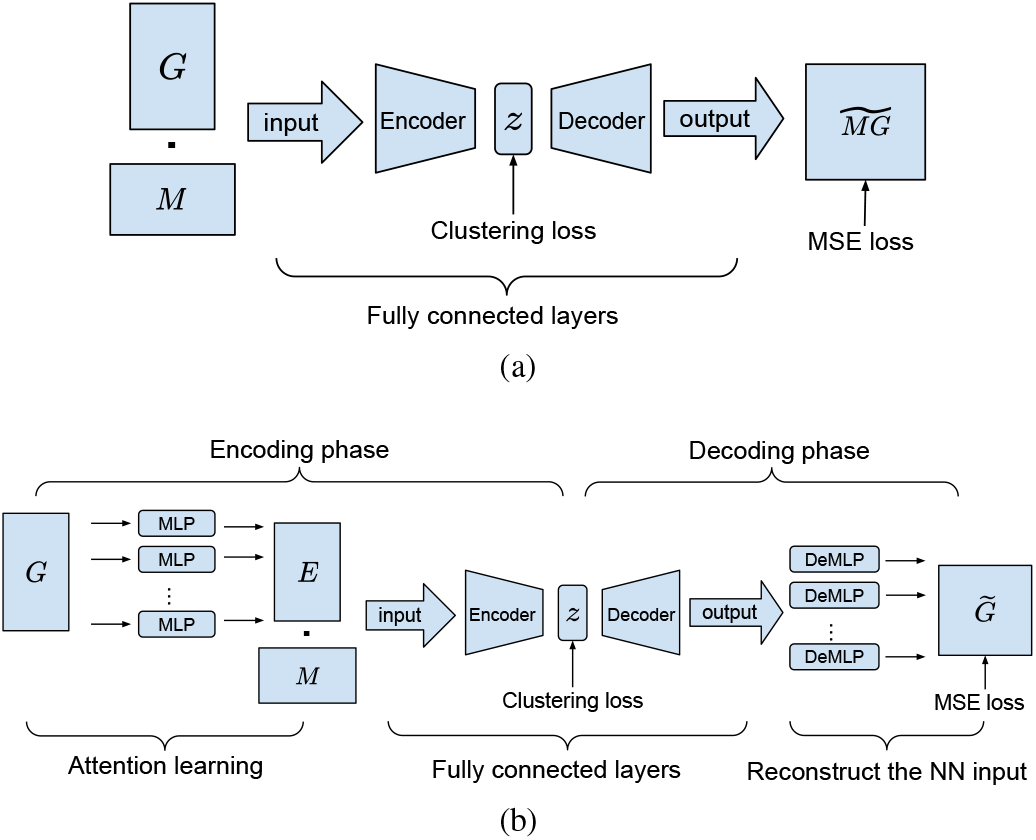
(a) is the pipeline without attention learning compared with the main pipeline, using pre-trained GloVe embedding directly as model input. (b) has an extra ‘deMLP’ stage comparing with the main pipeline. The ‘deMLP’ stage mirrors the MLPs in attention learning by trying to keep the pipeline symmetric. The idea is to construct a larger AE by extending the encoding and decoding phases.

**Fig. I.2.**
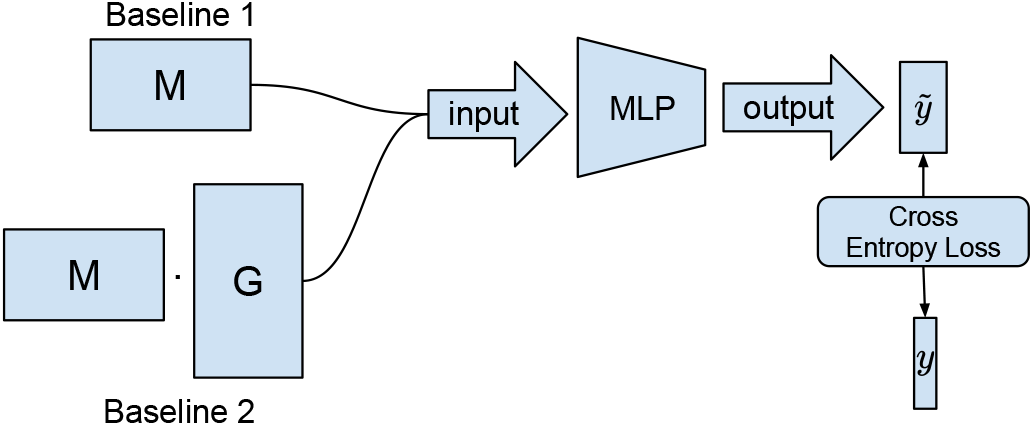
Two baseline classification pipelines. Baseline 1 uses pretrained GloVe embeddings mapped with patient-ICD encoding matrix as predictors; Baseline 2 uses patient-ICD encoding matrix only as predictors. The rest of the model stays the same with the main pipeline.

## Appendix II

### Training Details

We trained GloVe to produce 128-dimensional embeddings, with Xavier initialisation on the weights, Adam [13] as the optimiser and with batch size of 128. We also applied a learning rate scheduler (‘ReduceLROnPlateau’ in Pytorch): the learning rate would reduce with a factor of 0.1 when the loss of the validation set does not decrease for two consecutive epochs. For the *AE pre-train* and *joint-train*, we deployed a three-layer stacked AE with layer widths [128, 64, 32]. We set the attention dimension the same as in [6], 200. Leaky Rectified Linear Unit [19] was applied as the activation function for all layers apart from the last layer in encoder and the first and last layers in decoder where Tanh activation function was used. This follows the training practice executed in [9]. We also applied an early stopping criterion (the validation loss is not reducing for 4 consecutive epochs) in the training process. We ran the *AE pre-trained* without the clustering loss and with the default parameters of Adam until convergence. We then reduced the starting learning rate

**Fig. II.3.**
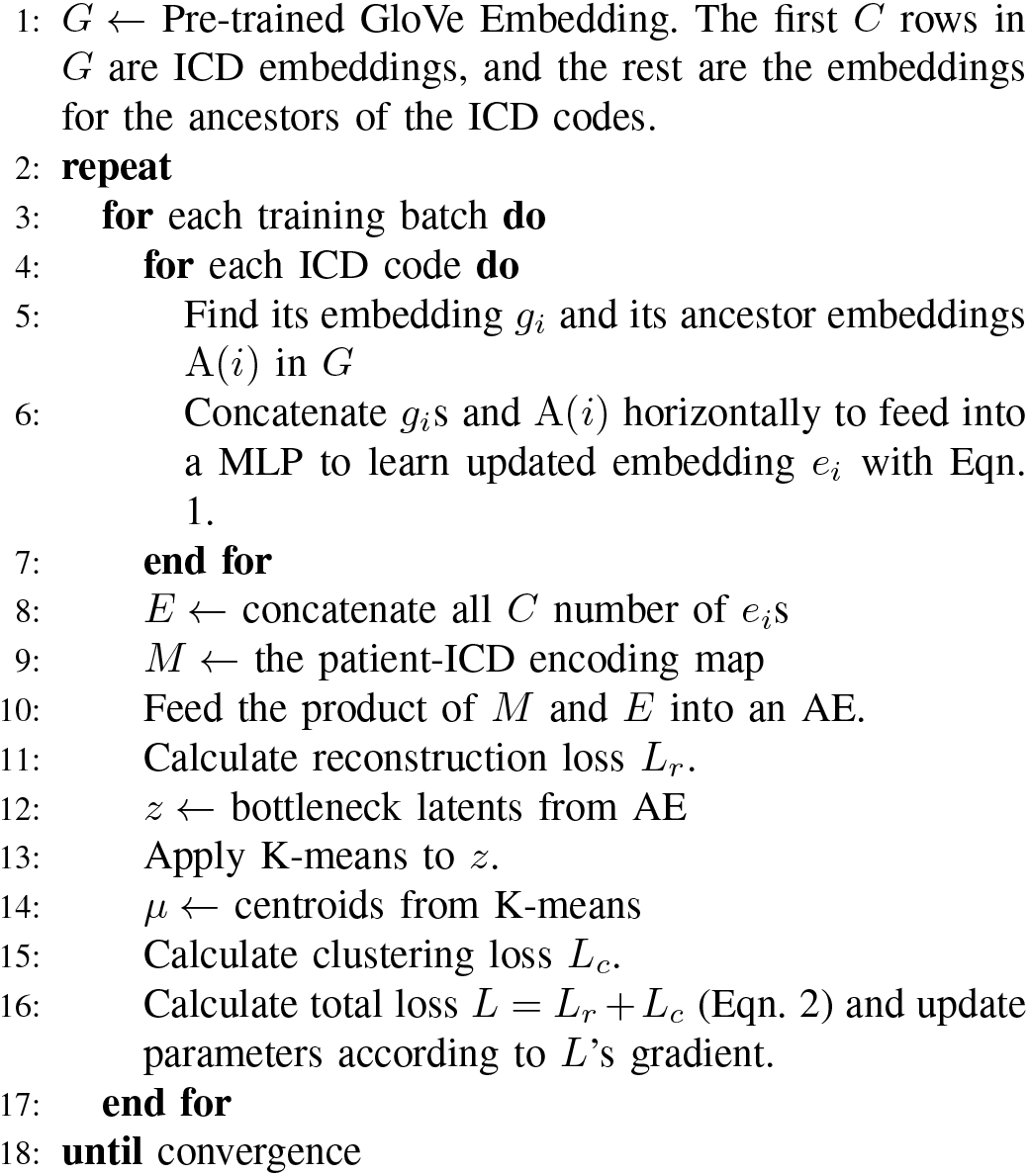
Algorithm of the end-to-end training in Fig. 2 B.

of Adam to 1*e* – 5, and further ran the *joint-train* with the joint loss function until convergence (this procedure is illustrated in Appendix Fig. II.3).

We did the model training on a 2.6 GHz 6-Core Intel Core i7 processor. The GloVe pre-train trains around 20,000 samples per second which takes roughly 2.5 seconds for one epoch for the 3266 ICD codes; both the AE pre-train and joint-train take roughly 15 seconds for one epoch for 2216 patients.

#### A. Computational Complexity

Based on Box A in Fig. 2, the computational complexity of the five main components is as follows: the embedding pretrain makes use of GloVe which has a worst-case complexity of 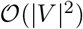 [29], where |*V*| is the vocabulary size, in our case, the number of unique codes; the graph attention has complexity of 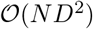 where *D* is the embedding dimensionality and *N* is the sample size; for autoencoder, given an architecture with fixed depth and layer widths, both training stages, without (pre-train) and with (joint-train) K-means loss have the complexity of 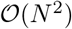; UMAP, by the authors of the original paper [21], has an empirical complexity of approximately 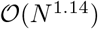; finally, for K-means, the complexity of Lloyd’s algorithm which was the algorithm adopted in our pipeline is 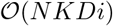, where *K* is the number of clusters and *i* is the number of iterations needed until convergence. Given a fixed setting (the maximum number of iterations set and *K* fixed), the complexity is 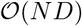 which is generally smaller than 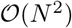. Therefore, the overall complexity of our pipeline would be 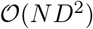.

## Appendix III

### Illustration for Over-training the Joint-train Stage

Fig. III.4 is an UMAP illustration for over-training at the joint-train stage.

**Fig. III.4.**
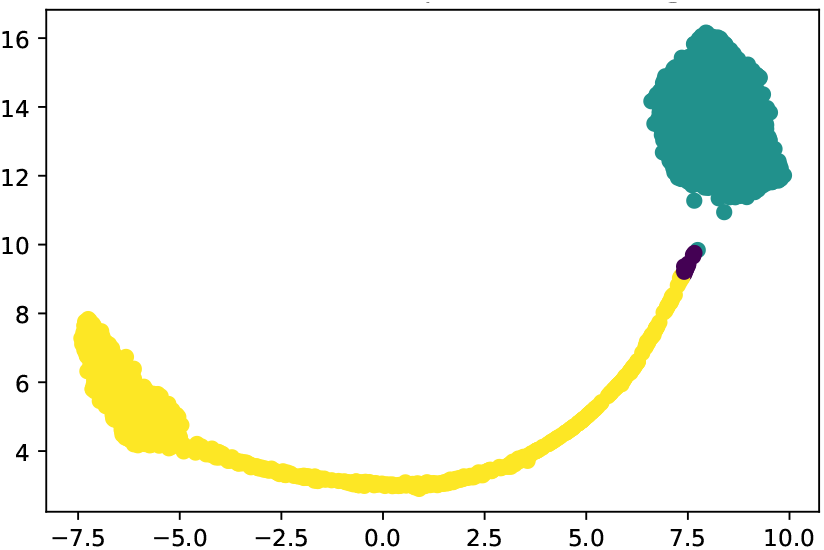
UMAP visualisations for the bottleneck latents after joint-train and for *k* = 2 (trained with two clusters). The latents are extract from applying the converged model to all of the OF patients. The difference between this figure and the ones shown in Fig. 8 is that we trained this model for more epochs and fine-turned the model for a longer time period. We observe that the yellow cluster is being stretched which results in the clustering algorithm (HDBSCAN) identifying some outliers at the end of the cluster tail.

## Appendix IV

### Patient Clustering Results

Figs. IV.5 and IV.6 show the top 10 CCS single-category and top 20 ICD code-level results for *k* = 2 respectively. Fig. IV.7 shows the results for *k* = 3 and *k* = 4.

**Fig. IV.5.**
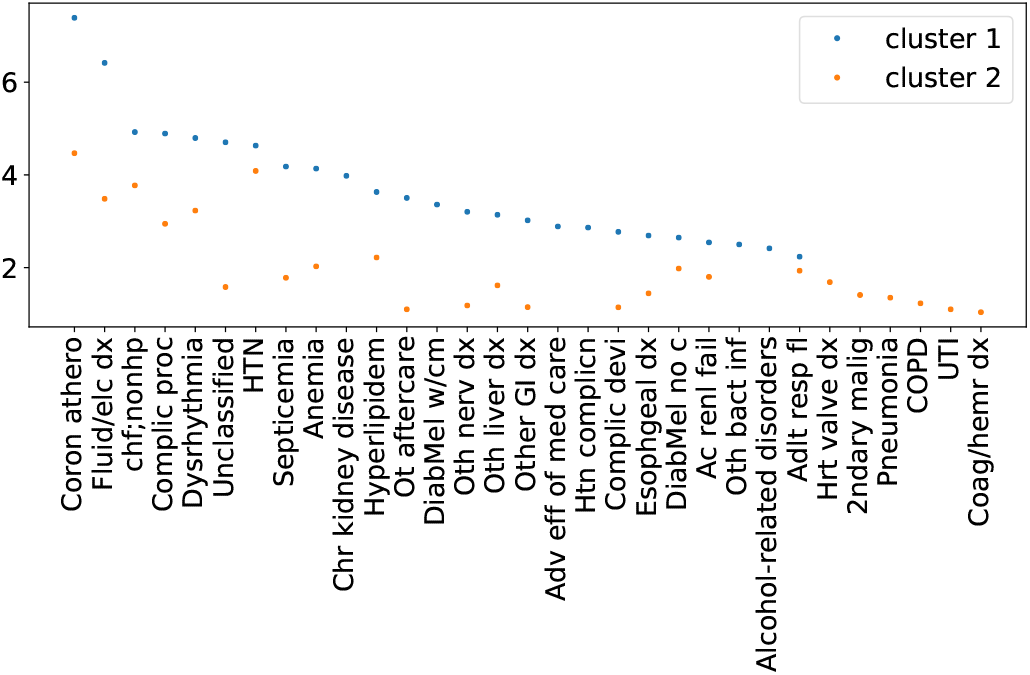
CCS single-level categories with the top 10 percent occurrence in the two clusters. The occurrence is normalised by cluster size and ordered in decreasing orders. The label ticks are abbreviated category names. The full category names can be found in Appendix V. Compared with Fig. 9, there are more overlapping categories between the two clusters.

**Fig. IV.6.**
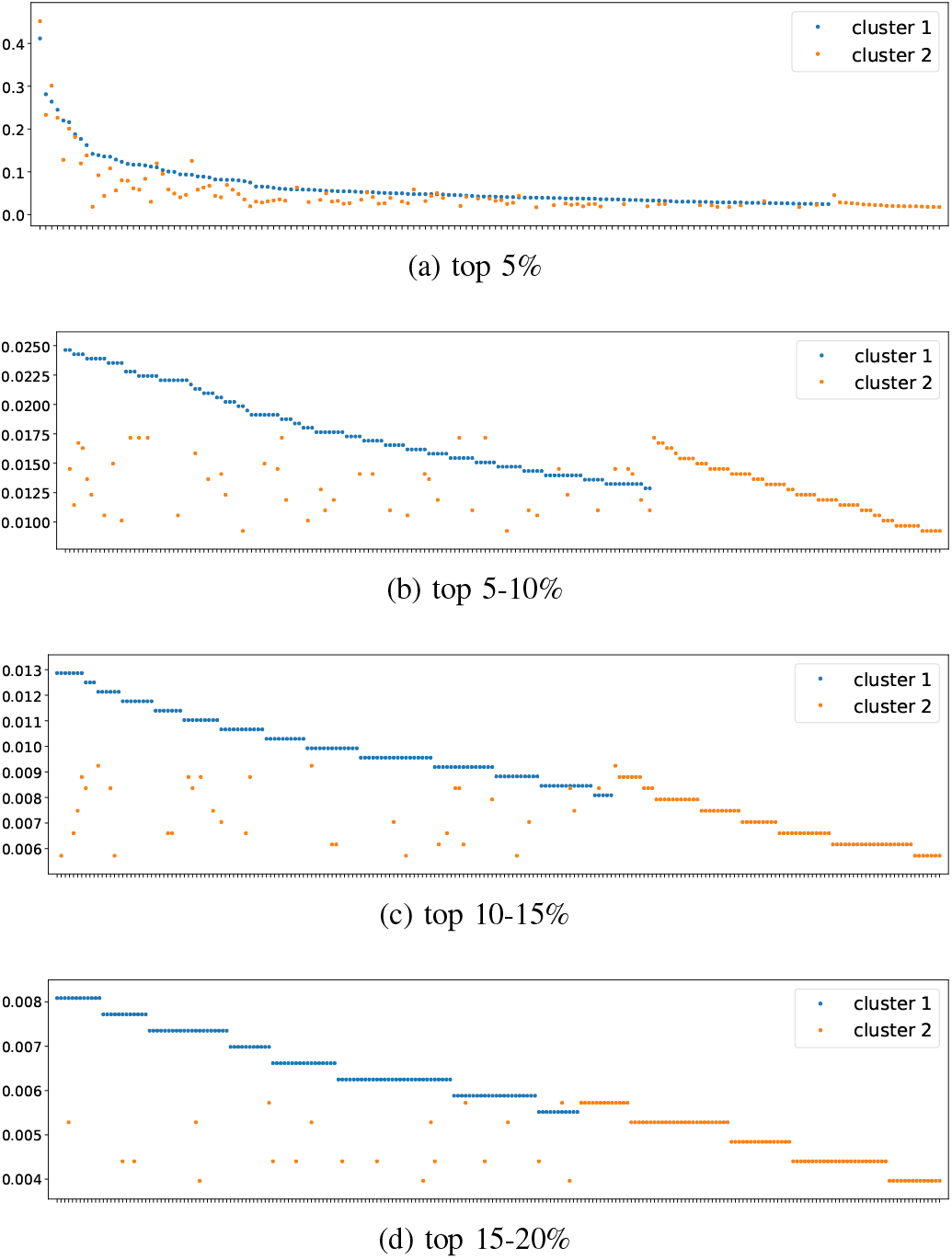
The pattern of the top 20 most frequently occurring ICD codes between the two clusters. Due to the vast number of ICD codes, the x-axis tick labels are omitted and the visualisation is split into 4 ranges, top 5% (a), top 5-10% (b), top 10-15% (c) and top 15-20% (d). The occurrence is normalised by cluster size and ordered in decreasing orders. The label ticks are ICD codes. Although it is hard to interpret by each ICD code, it is clear to see that the number of overlapping codes between the two clusters decreases from (a) to (d). It is clear to observe from the top to the bottom subfigures that the number of overlapping codes is decreasing and we are able to distinguish distinct codes between the clusters.

**Fig. IV.7.**
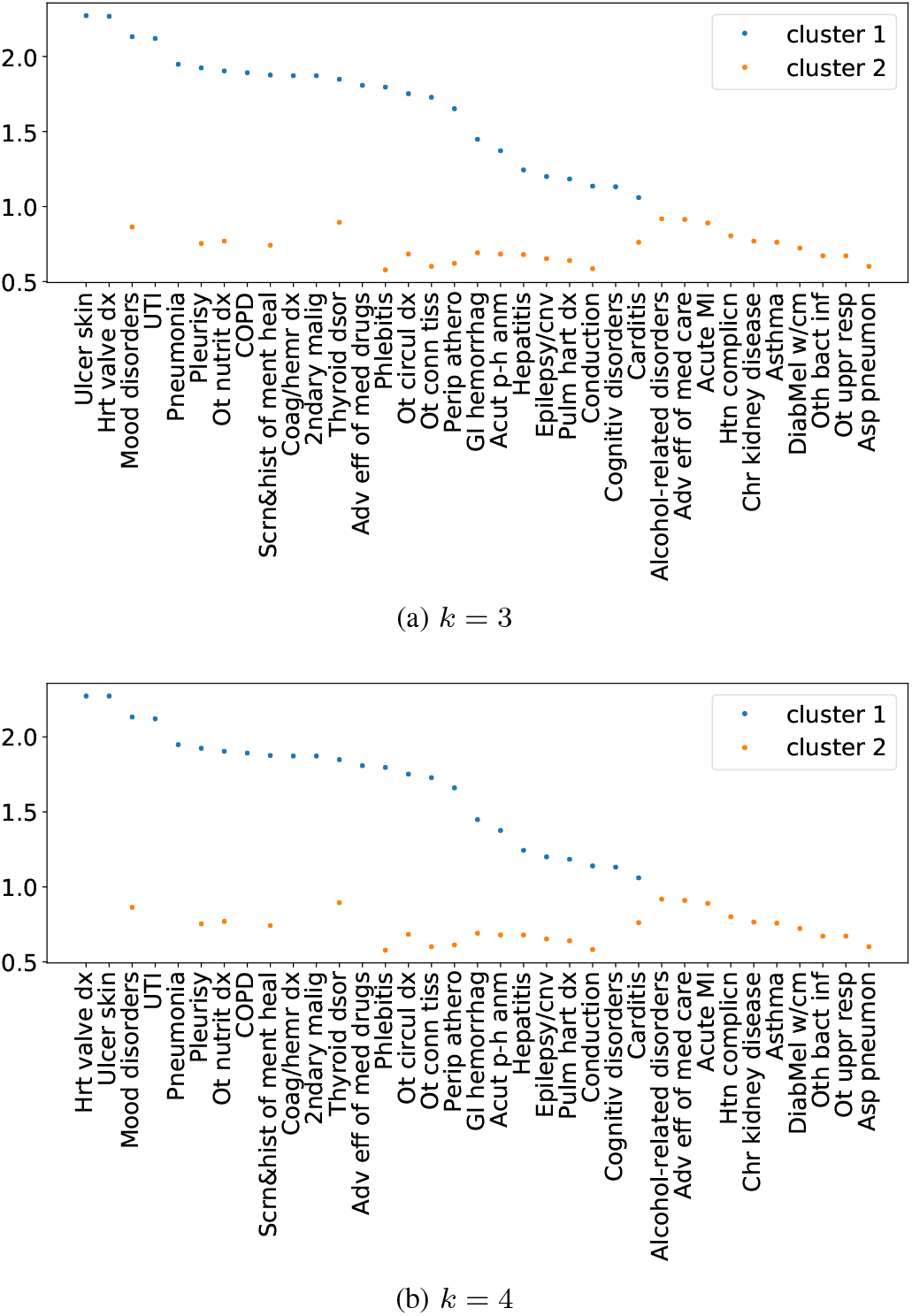
Top 10-20 most-occurred CCS single-level categories for cases of *k* = 3 (a) and *k* = 4 (b) in K-means. The occurrence is normalised by cluster size and ordered in decreasing orders. The label ticks are abbreviated category names.

## Appendix V

### CCS Single-level Category Lookup tables

**Fig. V.8.**
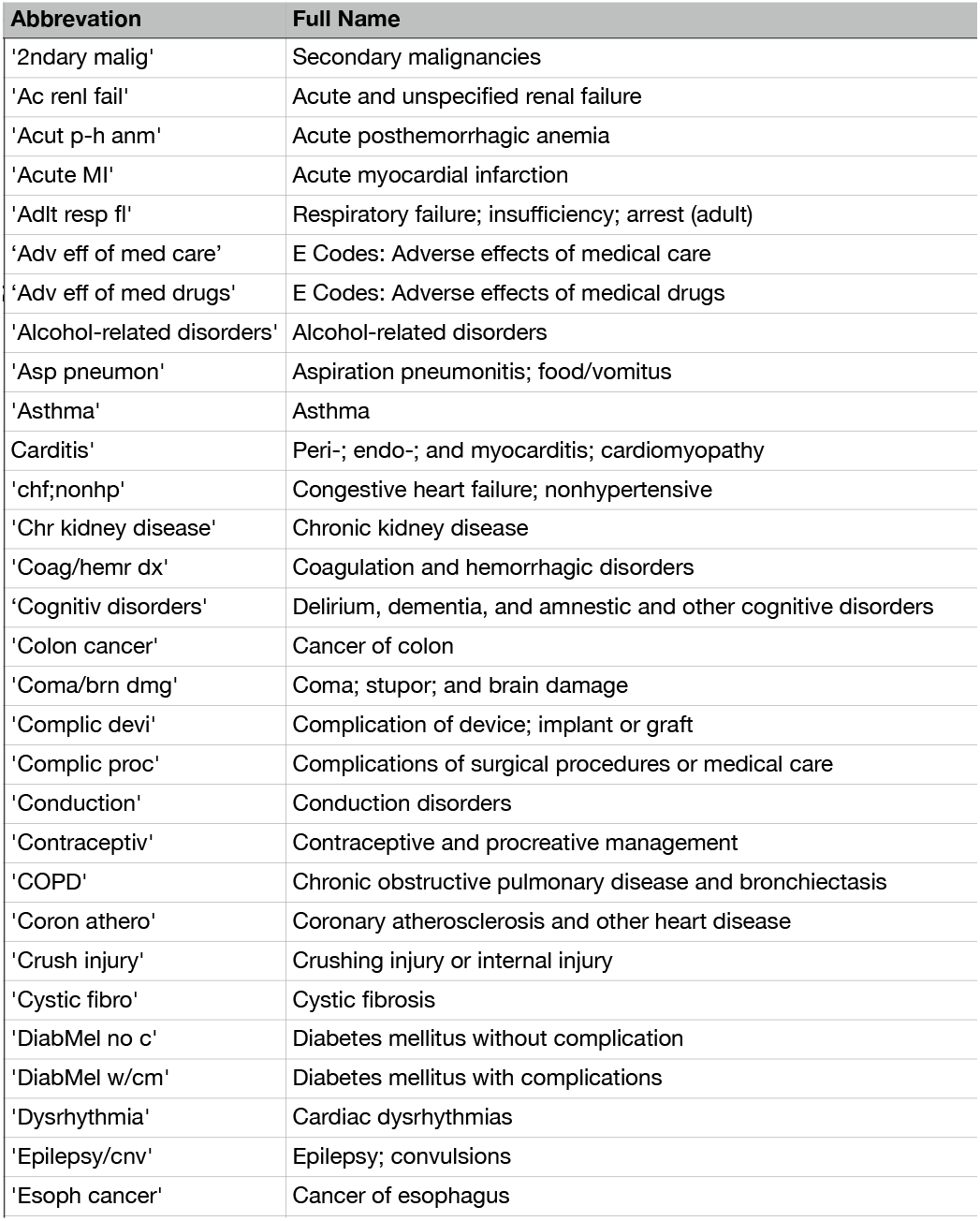
The first lookup table for the abbreviated CCS single-leve categories and their full category names.

**Fig. V.9.**
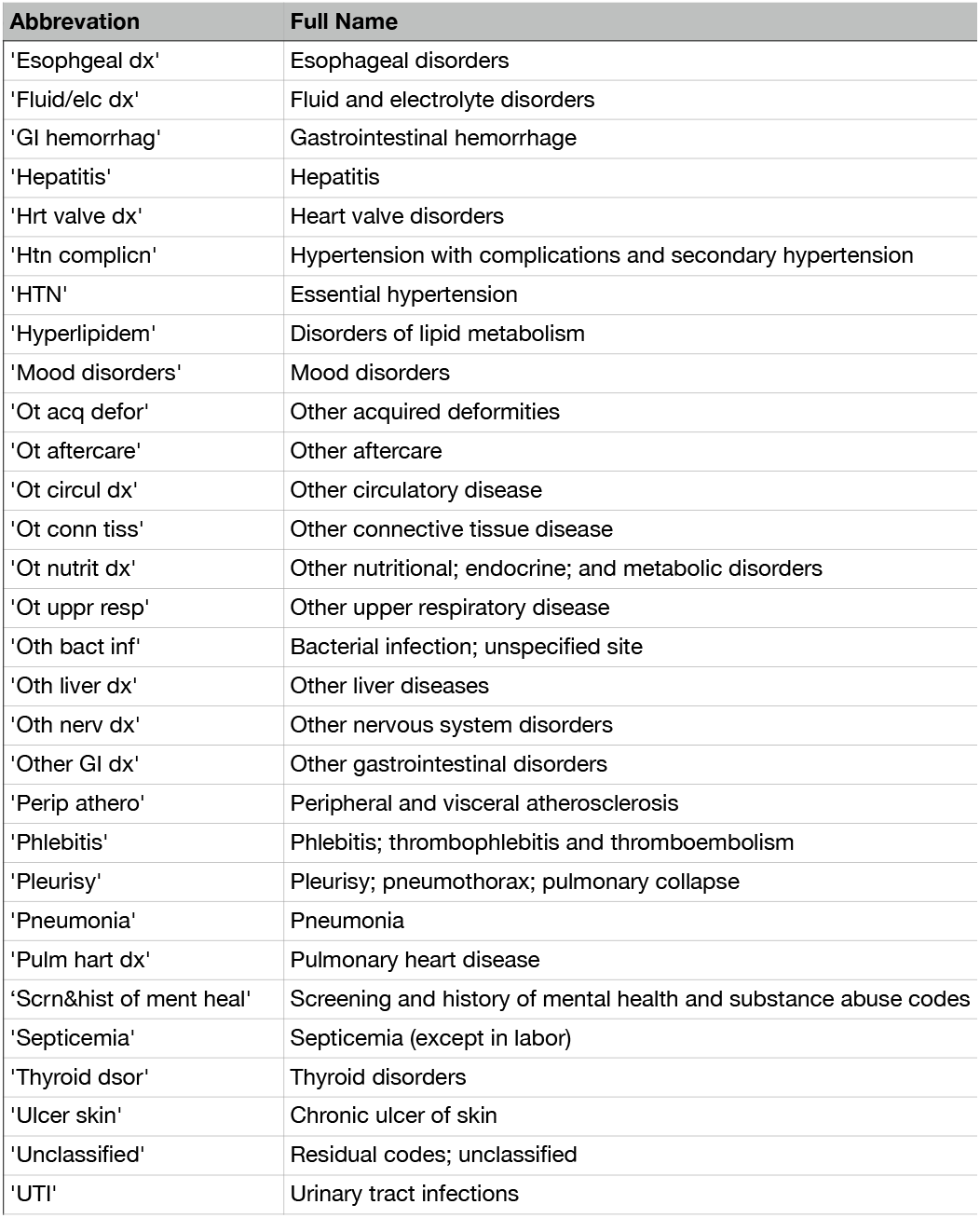
The second lookup table for the abbreviated CCS single-level categories and their full category names.

## Footnotes

1 the Github link will be attached upon acceptance.

## Notes

### Competing Interest Statement

The authors have declared no competing interest.

